# Torsional stress generated by ADF/cofilin on cross-linked actin filaments boosts their severing

**DOI:** 10.1101/380113

**Authors:** Hugo Wioland, Antoine Jégou, Guillaume Romet-Lemonne

## Abstract

Proteins of the Actin Depolymerizing Factor (ADF)/cofilin family are the central regulators of actin filament disassembly. A key function of ADF/cofilin is to sever actin filaments. However, how it does so in a physiological context, where filaments are interconnected and under mechanical stress, remains unclear. Here, we monitor and quantify the action of ADF/cofilin in different mechanical situations by using single molecule, single filament, and filament network techniques, coupled to microfluidics. We find that local curvature favors severing, while tension surprisingly has no effect on either cofilin binding or severing. Remarkably, we observe that filament segments that are held between two anchoring points, thereby constraining their twist, experience a mechanical torque upon cofilin binding. We find that this ADF/cofilin-induced torque does not hinder ADF/cofilin binding, but dramatically enhances severing. A simple model, which faithfully recapitulates our experimental observations, indicates that the ADF/cofilin-induced torque increases the severing rate constant 100-fold. A consequence of this mechanism, which we verify experimentally, is that cross-linked filament networks are severed by cofilin far more efficiently than non-connected filaments. We propose that this mechano-chemical mechanism is critical to boost ADF/cofilin’s ability to sever highly connected filament networks in cells.

---

A number of essential cellular processes rely on the regulated assembly and disassembly of actin filament networks (1, 2). The main proteins responsible for actin filament (F-actin) disassembly are the members of the Actin Depolymerizing Factor (ADF)/cofilin protein family (3–5). ADF/cofilin binds to ADP-F-actin in a cooperative manner, leading to the formation of ADF/cofilin domains (6–10). These domains make filaments locally more flexible, for both bending and twisting (11–13), and shorten their helical pitch (without changing their length)(14–16). Filaments consequently sever at, or near, domain boundaries (8–10, 17). ADF/cofilin-saturated filament fragments do not sever, since they contain no domain boundaries, but they efficiently depolymerize from both ends (10). In particular, ADF/cofilin-decorated filaments have barbed ends that can hardly elongate or get capped, and thus depolymerize extensively, even in the presence of monomeric actin or capping proteins (10).

We have recently measured the rate constants of these different binding, severing, and depolymerizing reactions (10). These results were obtained, as for many *in vitro* characterizations, by monitoring filaments that were barely constrained mechanically. In contrast, most filaments in cells are part of interconnected, or cross-linked, networks, and are exposed to various mechanical stresses. The specific activity of ADF/cofilin in this context is unclear.

Mechanical stress has long been proposed to potentially enhance severing by cofilin (18, 19). Filaments immobilized on coverslips were reported to sever preferentially in bent regions when exposed to actophorin, a member of the ADF/cofilin family found in amoeba (18). Tension has been reported to protect filaments from ADF/cofilin binding and severing (20) and so has, very recently, formin-induced filament torsion (21). A recent theoretical study proposes that buckled filaments are easier to sever, while twisting a filament would mostly favor the dissociation of cofilin (22).

In addition to the external application of mechanical stress, seemingly passive mechanical constraints such as filament anchoring may also play a role. For instance, it has been proposed that mechanical constraints could enhance the action of ADF/cofilin on filaments anchored to a coverslip surface (23). In cells as well as in vitro, filaments cross-linked into bundles by fascin have been reported to sever faster than individual filaments when exposed to ADF/cofilin, and several explanations have been proposed, including a contribution of mechanical constraints (24).

A primary aspect is that, according to structural data, ADF/cofilin domains locally change the helical pitch of actin filaments (14–16). When filaments are anchored or cross-linked, their overall twist is constrained and this feature thus appears to be in conflict with ADF/cofilin binding. Existing data thus indicate that twist constraints and torque are likely to be key parameters affecting cofilin activity. Whether they contribute to favor or hinder cofilin binding, and/or severing, and to what extent, are all open questions.

Here, we investigate how ADF/cofilin binding and severing are affected by mechanical tension, by bending, and by constraints applied on the filament’s twist. We show that cofilin generates a torsional stress when binding to twist-constrained filaments, leading to a drastic enhancement of their severing.

## Results and Discussion

### Cofilin binding induces a torsional stress on actin filaments which cannot freely rotate around their main axis

To directly assess the effect of ADF/cofilin binding on filament torsion, we monitored the polarization of the light emitted by single labeled actin subunits (25, 26), incorporated within filaments that were anchored by either one or two ends in a microfluidics chamber (Fig. 1). In the absence of cofilin, the polarization index of labeled subunits fluctuated mildly around a constant value, indicating that these subunits remained pointing in a fixed direction. When exposed to cofilin, the polarization index began to vary, reflecting the rotation of the subunits’ orientation around the filament’s main axis (for 13 out of 15, and 12 out of 25 observed subunits on filaments anchored by one, or two ends, respectively. Fig. 1C and Fig. 1D show subsets of 4 representative measurements for each condition). Variations of the polarization index were more pronounced and more regular when only one filament end was anchored (Fig. 1 and Fig. S1E). These observations are in agreement with numerical simulations that we performed (Fig. S1) assuming that cofilin-decorated regions have a 25% shorter helical pitch (14) and are 18-fold more compliant to twist than bare regions (11).

**Fig 1:**
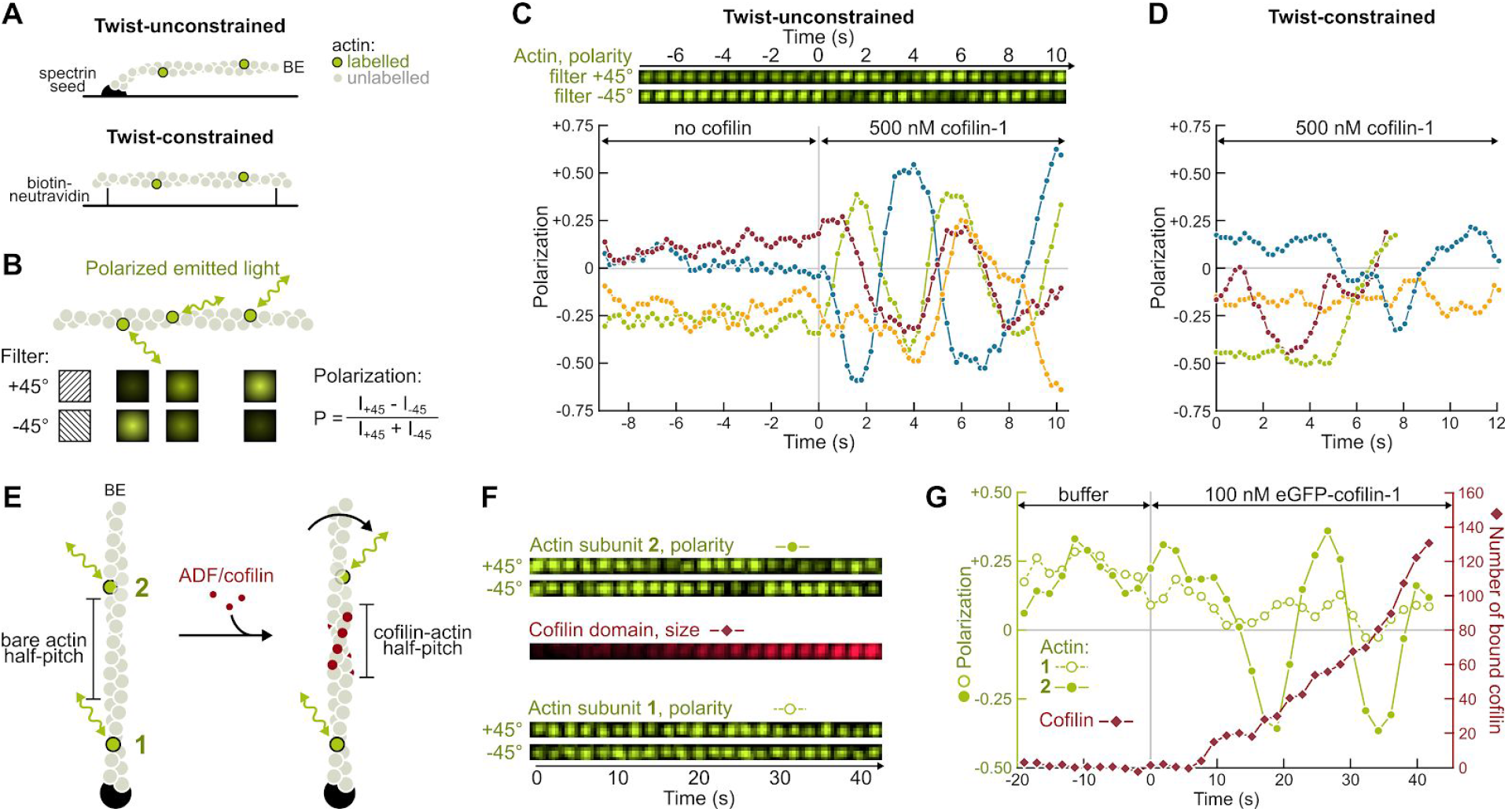
Direct visualization of the torsional stress induced by ADF/cofilin binding to anchored actin filaments. (A) In a microfluidics chamber, filaments with a low fraction of labeled subunits are either polymerized from surface-anchored spectrin-actin seeds (single anchor, twist is unconstrained), or anchored by their sides via biotin-neutravidin bonds (between two anchoring sites, the twist is constrained). (B) The polarization of the emitted light indicates the orientation of a single actin subunit. The polarization P = (I_+45_ - I_−45_)/(I_+45_ + I_−45_) is determined by measuring the emitted intensity through two different polarization filters (I_+45_ and I_−45_). (C) Top: for a twist-unconstrained filament, timelapse of the fluorescent intensities measured for a single actin subunit through the 2 polarization filters. Bottom: variation of the polarization P over time, measured for a single labeled subunit on 4 different twist-unconstrained filaments. The green curve corresponds to the same subunit as the timelapse shown above. 500 nM cofilin was injected from time t=0 onward. Here only, filaments were polymerized for 15 to 20 minutes but not aged further F-actin was partially in a ADP state to slow down cofilin binding (see Fig SX for full ADP-F-actin). (D) Variation of the polarization P over time, measured for a single labeled subunit on 4 different twist-constrained filaments, exposed to 500 nM cofilin from time t=0 onward. (E) Sketched from above: ADF/cofilin binding shortens the helical pitch and thus rotates the filament segment located between the ADF/cofilin domain and the free barbed end (BE), including subunit 2, while subunit 1’s orientation does not change. (F) Timelapses of the fluorescent intensities measured in the configuration sketched in (E): the two labeled actin subunits, each seen through the 2 polarization filters, as well as the total intensity of the growing eGFP-cofilin-1 domain positioned between these two labeled subunits. (G) Polarization of the same two labeled actin subunits, compared with the estimated number of cofilin monomers bound between these two subunits. 250 nM eGFP-cofilin-1 was injected from time t=0 onward. On this specific example, subunit 2 (filled green symbols) made one full rotation at t≈30s, when approximately 60 eGFP-cofilin-1 molecules had bound the filament.

Filament rotation induced by cofilin binding could be most clearly characterized by monitoring the appearance and growth of a labeled cofilin-1 domain, between the anchored filament end and a labeled actin subunit (Fig. 1E-G). Within the resolution of our experiment, this subunit began to rotate when the fluorescent signal from the ADF/cofilin domain was first detected between the anchoring point and the subunit. We have calibrated the fluorescence intensity of eGFP-cofilin-1, and could thus estimate that one full turn of the filament was achieved when 60-120 cofilin molecules were added (Fig. 1G, observed on 4 different filaments). This number is close to what one would deduce from the reported reduction in helical pitch for cofilin-decorated filaments, which leads to an estimated 80 cofilins to cause one full turn. Consistently, we found the rotation velocity of the filaments to be correlated with cofilin concentration, which modulates domain nucleation and growth rate (Fig. S2).

These results show that filaments with a free end rotate around their main axis upon cofilin binding, thereby preventing the application of torsional stress. In contrast, cofilin domains decorating a filament segment between two anchoring points will impose a mechanical torque on this segment: both bare and decorated regions will be under-twisted relative to their spontaneous helicity (Fig. S1B).

### Twist-constrained filaments are severed more efficiently by ADF/cofilin

We next sought to examine the consequences of this mechanical torque. To do so, we compared the action of cofilin on filaments held between two anchoring points (i.e. twist-constrained, thus experiencing a torque as cofilin binds) with its action on filaments held by a single anchoring point (i.e. free to rotate, and thus not subjected to torque). These two configurations were achieved simultaneously in the same microfluidics chamber, by anchoring sparsely biotinylated filaments with one flow direction and then exposing them to labeled cofilin-1 with an orthogonal flow direction (Fig. 2A). We monitored the increase in the fluorescence signal of eGFP-cofilin-1 on each population and found that cofilin binds equally well to twist-unconstrained or twist-constrained filaments (Fig. 2C).

**Fig 2:**
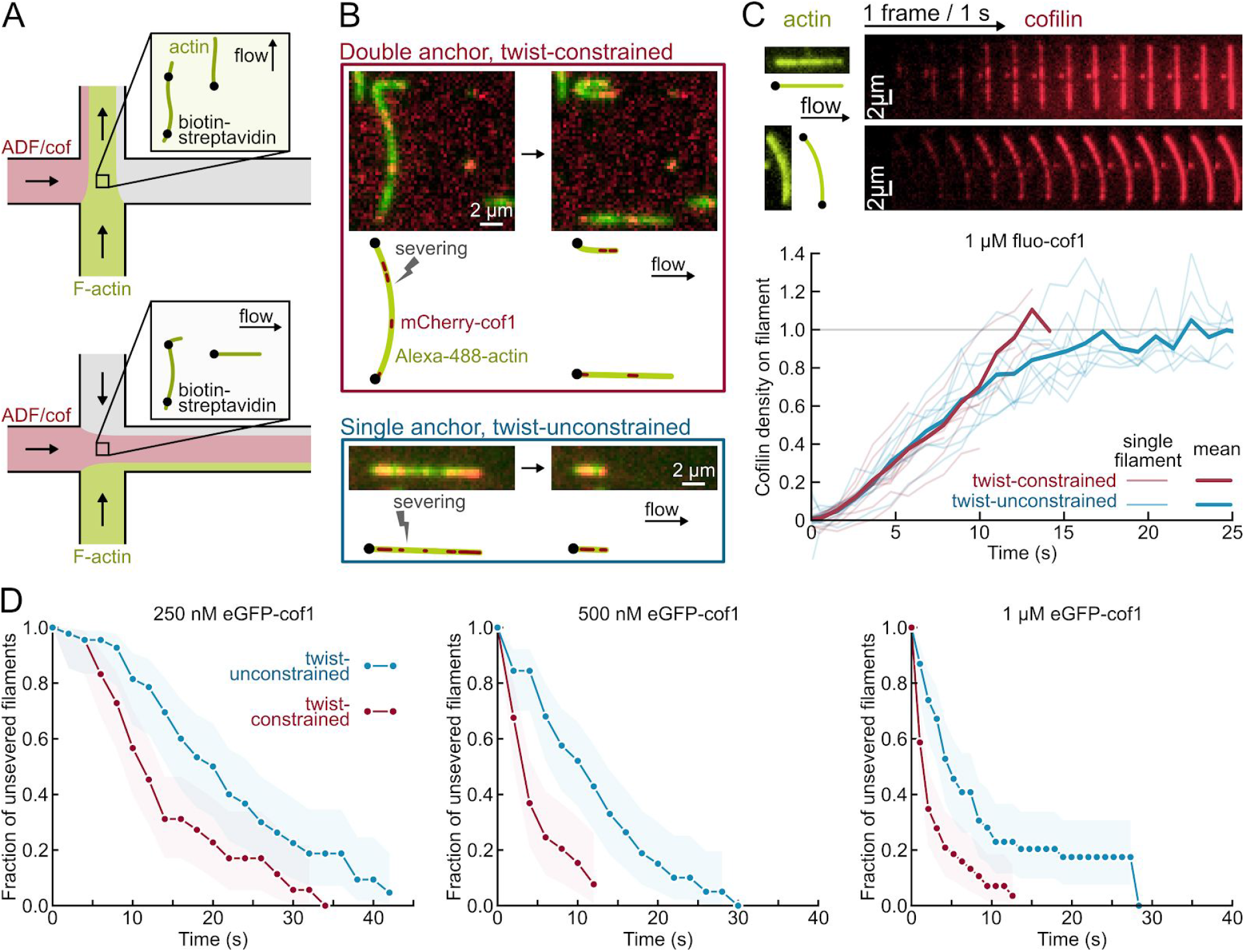
Constraining the twist of actin filaments accelerates their fragmentation by cofilin. (A) Experimental setup, seen from above. Sparsely biotinylated F-actin is injected across the chamber and binds neutravidin-biotin-PLL-PEG. A perpendicular flow (containing ADF/cofilin) reveals the position of anchored points along the filament. (B) Examples of severing events for actin segments anchored at one (blue, bottom) or both ends (red, top). Severing occurs at cofilin-1 domain borders. See also Movie 1. (C) Cofilin-1 binding is unaffected by twist constraint. Top: raw data for two filaments, with unconstrained and constrained twist. Bottom: mean cofilin density averaged over ten filaments for each condition. Condition: 1 μM eGFP-cofilin-1 from time t=0 onward. Images acquired using TIRF. (D) Actin segments with constrained twist fragment faster. Fraction of unsevered actin segments for twist-constrained and unconstrained, exposed to different cofilin concentrations from time t=0 onward. The survival fraction is calculated over 49 filaments for each condition, blindly selected with the same length distributions (SI Methods) and <L> = 6.6 ± 1.6 μm (std). Shadows represent 95% confidence intervals.

We measured the survival fraction of unsevered filaments in each population and found that twist-constrained filaments were severed significantly faster (Fig. 2D). Severing occurred near domain boundaries, both on unconstrained and constrained filaments (Fig. 2B). In the absence of cofilin, no significant severing was observed in either population (Fig. S3). We also verified that, after a severing event on a twist-constrained filament, the two resulting single-anchored filament fragments exhibited the same, lower severing rate as filaments in the twist-unconstrained population (Fig. S4). The enhanced severing was also observed with ADF or at pH 7.0 (Fig. S5).

It thus appears that cofilin severing is enhanced by the torsional stress induced by cofilin binding to filament segments between two anchoring points.

### Filament bending enhances severing by cofilin, while tension has no effect

We next examined if an externally applied stress could alter the severing rate. Due to the helical nature of the actin filament, twist and bending are coupled (27) and we thus expected filament bending to also enhance severing by cofilin. To test this idea, we anchored short phalloidin-stabilized filaments to the bottom of the flow chamber and, thanks to the flow, we imposed a different direction to the unanchored filaments that elongated from them (Fig. 3A). We found that larger angular differences between the anchored and free segments, which correspond to higher local curvatures near the anchored segment, led to faster severing by cofilin in that region (Fig. 3A-C).

**Fig 3:**
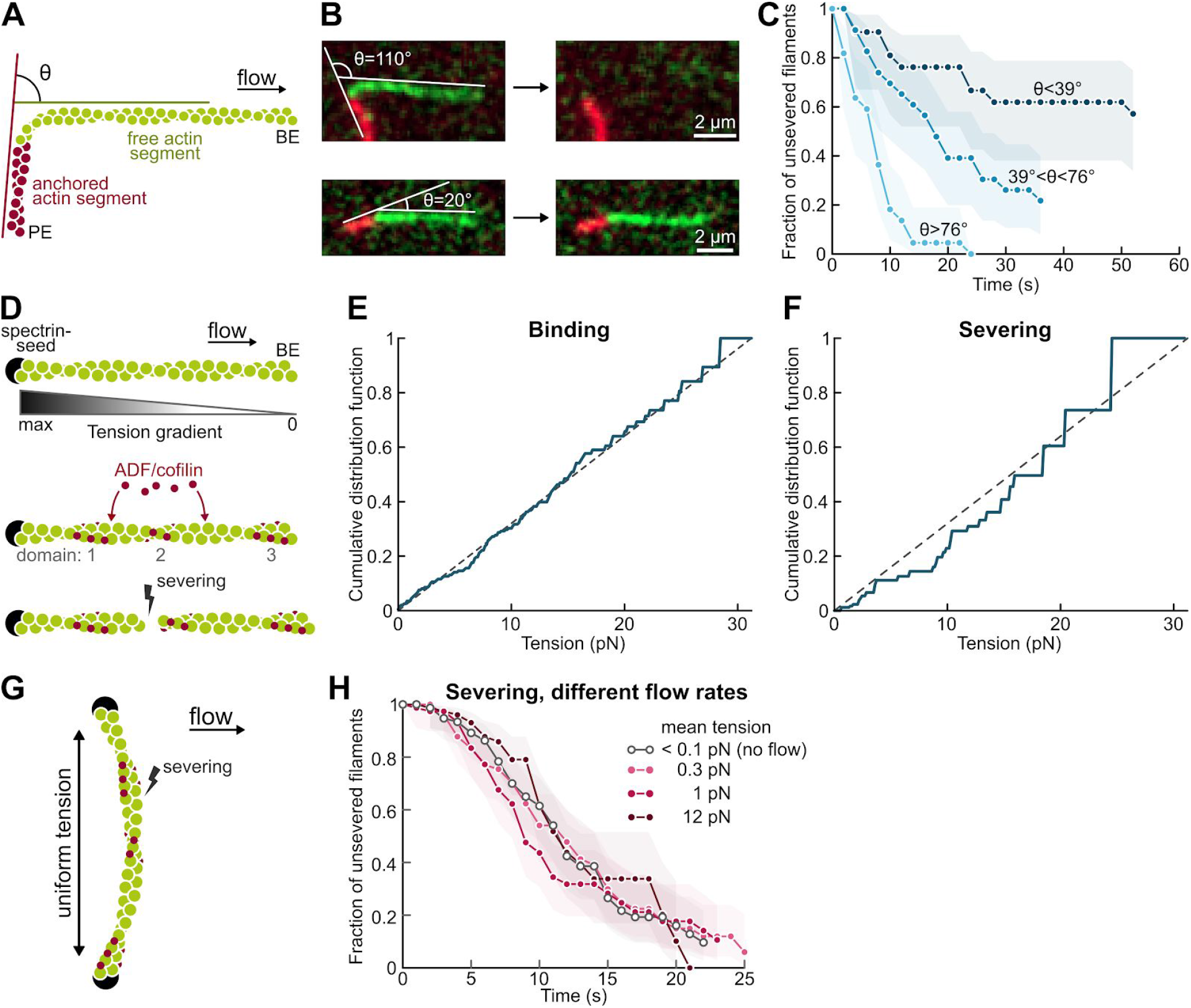
ADF-cofilin-induced severing is also accelerated by bending but not by tension. (A) To assess the effect of curvature, a segment of biotin-F-actin is stabilised with rhodamine-phalloidin and left to bind the neutravidin-biotin-PLL-PEG surface with a random orientation (red). A second segment, that does not bind the surface, is then polymerized from this seed (green). Only severing events taking place in the curved green region were considered. (B) Example of two filaments. The angle θ is defined as the angle between the flow direction and direction of the anchored segment. (C) ADF/cofilin-induced severing is stronger on curved filaments. The filament population is split into 3 subsets of equal size and increasing angle θ. Conditions: 2 μM ADF, injected from time t=0. N = 21, 23, 22 from low to high angles. L = 5.5 ± 0.7 μm, the tension applied in the curved region was 0.2 ± 0.03 pN. Shadows represent 95% confidence intervals (SI Methods). (D). The viscous drag of the fluid on a filament anchored by only one end generates a linear tension gradient. (E-F). The cumulative distribution of cofilin domain appearance (E) and severing events (F) on single-anchored filaments follow a linear function, indicating that both events are uniformly distributed and thus do not depend on the local tension. For (E), 42 pointed end-anchored filaments were exposed to 0.4 μM mCherry-cofilin-1 for 8 sec, and 117 domains were located on them. For (F), barbed end-anchored filaments were exposed to 1 μM mCherry-cofilin-1 for 5 sec and 31 severing events were spotted. (G). Anchoring actin filaments by both ends results in a nearly uniform tension, proportional to the filament length and the flow rate. (H). Survival fraction, with respect to severing, of double-anchored filaments exposed to different flow rates and thus to different tensions. 300 nM eGFP-cofilin-1 was continuously injected from time t=0 onward. All four populations where blindly selected with the same length distributions, <L> = 4.9 ± 1.2 μm (N = 76 for each flow rate).

The flowing solutions also put filaments under tension. When filaments are anchored by a single point, they are exposed to a linear tension gradient (28) while filaments anchored between two points, perpendicular to the flow, are exposed to a nearly uniform tension (Supp Text). We have modulated the tension (up to 30 pN) by varying the flow rates, and found that it had no effect on the binding nor the severing activities of cofilin: domain nucleation events and severing events occured homogeneously over filaments exposed to a gradient of tension (Fig. 3D-F) and twist-constrained filaments severed with the same rate, independently of the applied tension (Fig. 3G-H). In addition, severing events were homogeneously distributed over twist-constrained filaments, anchored by their two ends (Fig. S6). Since these results differ from previously published observations (20) we have repeated these experiments with different isoforms, different force ranges, and different anchoring strategies (Fig. S7). They all confirmed that, in our experiments, filament tension had no effect on ADF/cofilin activity. Report of a mechanosensitive disassembly of filaments in cells (29) thus indicates that filament tension probably affects other factors modulating the activity of cofilin, such as tropomyosins (30).

### A simple model accounts for the torque-induced enhancement of severing

To further describe and quantify the enhanced severing of twist-constrained filaments by ADF/cofilin, we have recapitulated our results in the following model (summarized in Fig. 4A, and detailed in Supp Text), which we compared to our experimental data thanks to numerical simulations. To account for ADF/cofilin cooperative binding, we assume domain nucleation to follow a quadratic dependence on cofilin concentration, and grow with the rate constants that we have previously measured (10). When twist is constrained, ADF/cofilin domains nucleate and grow with the same rates as on twist-unconstrained filaments, as indicated by our observations (Fig. 2C). A simple energy balance also supports this hypothesis: we can estimate that the energy benefit of cofilin binding is much larger than its torque-induced energy cost (Supp Text). The growth of a cofilin domain applies a mechanical torque Γ on the double-anchored filament. Using published values of torsional stiffness for (stiffer) bare and (softer) cofilin-decorated F-actin (11), we can compute Γ, which is uniform throughout the filaments, as a function of the cofilin coverage ratio (Supp text). We find that, due to the greater flexibility of cofilin-decorated regions, this torque rapidly reaches its maximum value (Fig. 4B).

**Fig 4:**
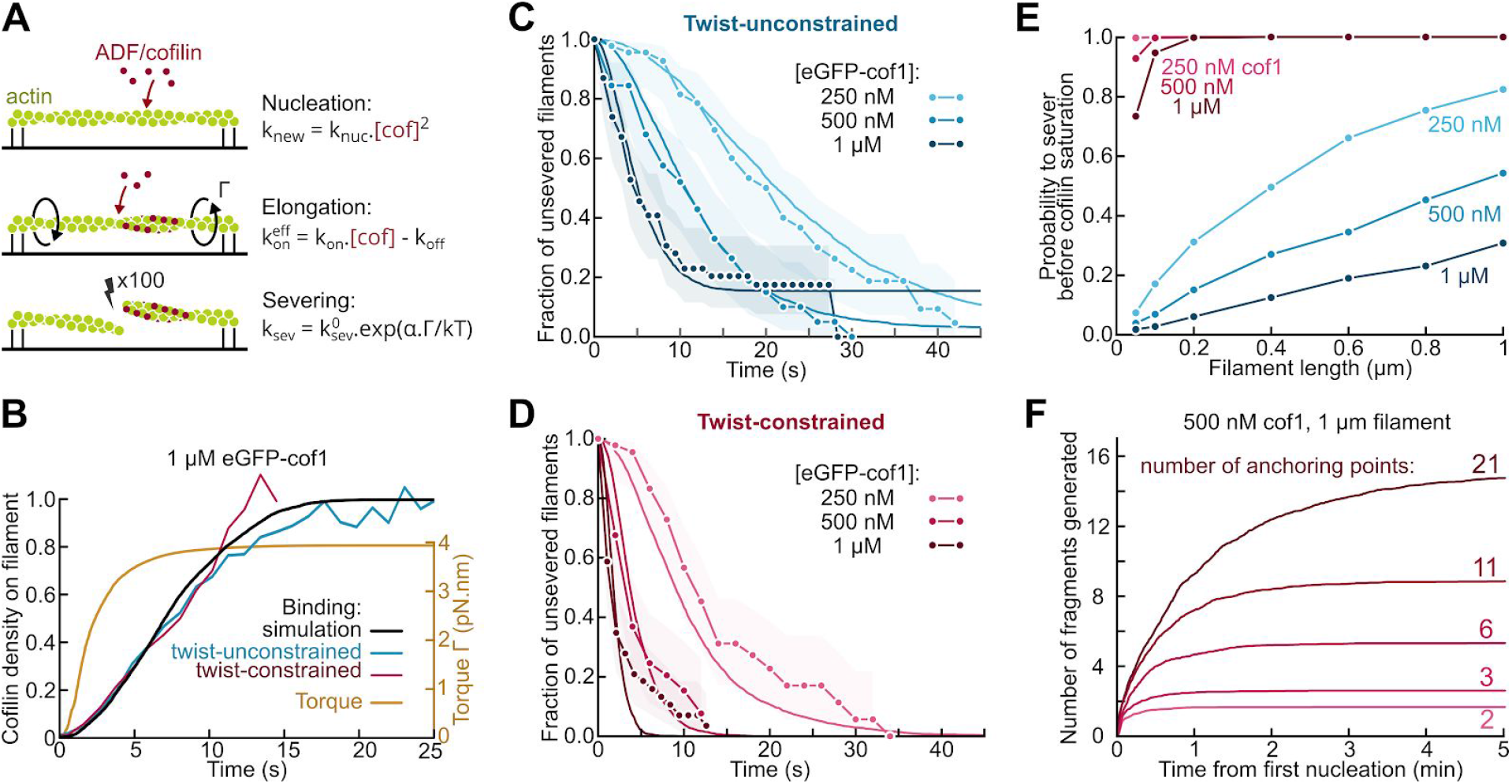
Model for the torque-induced enhancement of severing by cofilin. (A) Sketch and summary of the model used for simulations. (B) Computed cofilin density and resulting torque on twist-constrained filaments. The experimental curves for cofilin density (in red and blue) come from Fig. 2C. The maximum torque applied on twist-constrained filaments corresponds to an under-twist of approximately 5 rotations per micrometer. (C-D) Fit of the fraction of unsevered filaments in free (C) and constrained twist (D) conditions. k^0^_sev_ = 2.4*10^−2^ s^−1^ was determined by fitting the data at 1 μM cofilin with unconstrained twist, and α = 5 was then determined by fitting the data at 1 μM cofilin with constrained twist. All other curves were simulated using these same parameters, with no further adjustment. The experimental curves are from Fig. 2D. Simulations were performed over 50-fold larger samples of filaments with the same lengths. Shadows represent 95% confidence intervals (SI Methods). (E) Probability for a filament to sever before being saturated by cofilin-1 depending on its length and cofilin concentration, when twist is free (blue curves) or constrained (red curves). Each probability was computed by simulating 1000 filaments. (F) Simulated number of actin fragments generated by cofilin, on filaments anchored by at least their two ends plus additional points, randomly positioned along their length. Each curve shows the mean over 100 simulations. The curve reach a plateau when the filament is saturated by cofilin.

Severing occurs at domain boundaries. Fitting the survival fraction for twist-unconstrained filaments (Fig 4C) allowed us to determine the zero-torque severing rate constant k^0^_sev_. Since actin is partially labeled here, this severing rate constant is larger than the one we have measured in earlier work on unlabeled F-actin (10). We assumed that torque increased the severing rate exponentially, following a modified Bell model:

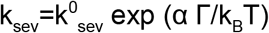

where α, quantifying the torque-sensitivity of severing, was the only unknown parameter and was determined by fitting the experimental survival fractions for twist-constrained filaments (Fig. 4D). Our model appears to be in very good agreement with our experimental data. We can estimate the maximum cofilin-induced torque to be approximately 3.9 pN.nm (Fig. 4B), based on published values of torsional stiffness, which are of the order of 10^−27^ N.m^2^ (11).

Recent computations (22) indicate that these numbers correspond to intersubunit torsional rigidities and that the filament torsional rigidity would be approximately 10-fold larger (31, 32). Our observation, using polarization microscopy, that individual subunits located micrometers away from the anchored end of the filament have a well-defined orientation (Fig 1), appears consistent with these larger values of torsional rigidity.

These values would lead to a larger estimate of the maximum torque and thus to a lower value of α, but our conclusion would remain: as cofilin domains nucleate and grow on twist-constrained filaments, they rapidly generate a torque, thereby enhancing the severing rate per cofilin domain. The severing rate is increased over 20-fold when 10% of the filament is decorated by cofilin, 50-fold when 20% is decorated, and up to 100-fold when the filament is nearly saturated by cofilin.

### Constraining a filament’s twist allows it to sever before being saturated by cofilin

Since severing occurs at the boundaries between cofilin domains and bare filament regions, cofilin-saturated filaments do not sever. Thus a factor that will determine the efficiency of severing is its ability to occur before the filament is fully decorated by cofilin. Enhancing severing with torsional stress not only allows it to occur faster, it may also allow it to happen on segments that would otherwise not sever at all. We expected this effect to be more pronounced on short filaments, which are more prone to become saturated without severing. This situation is certainly common in cells, where filament segments between crosslinks can be a few 100 nm long. However, individual severing events are difficult to resolve at such short length scales in our experiments.

Therefore, in order to investigate and quantify this point further, we have performed numerical simulations using our knowledge of the different reaction rates. We found that the torque-induced amplification of severing indeed allows cofilin to sever filaments in conditions where they would otherwise reach saturation without being severed (Fig. 4E). The difference is particularly strong for short filaments, which will be faster to saturate with cofilin. Note that, in this race against saturation, the enhancement of severing is made particularly effective by the fact that a significant torsional stress is already imposed by low densities of cofilin (Fig. 4B). Similarly, multiplying anchoring points allows cofilin to break filaments into more fragments (Fig. 4F).

This result explains why severing is more efficient when filaments are immobilized on a coverslip densely coated with myosins (23). As speculated by the authors of this work, cofilin binding generates a torsional stress which cannot relax when filaments are immobilized on a surface, leading to an enhanced severing rate. We show here that every filament segment between two anchoring points is likely to be severed before being saturated by cofilin, thanks to this torsional stress.

### Cross-linked networks sever much faster than unconnected filaments

Our results imply that a population of interconnected actin filaments, with a high density of cross-links, would sever much faster and much more than the equivalent non-connected filament population. In order to test this prediction, we have performed experiments on filament networks in T-shaped flow chambers (Fig. 5A). Preformed biotinylated actin filaments, with a 20:1 unlabeled-to-labeled filament ratio, were injected in the short end of the T-shaped chamber, which was then sealed, thereby creating a dead end which contained the filaments. We performed similar observations where all the filaments were fluorescently labeled, but having only a fraction of labeled filaments allows one to monitor and quantify single events (33).

**Fig 5:**
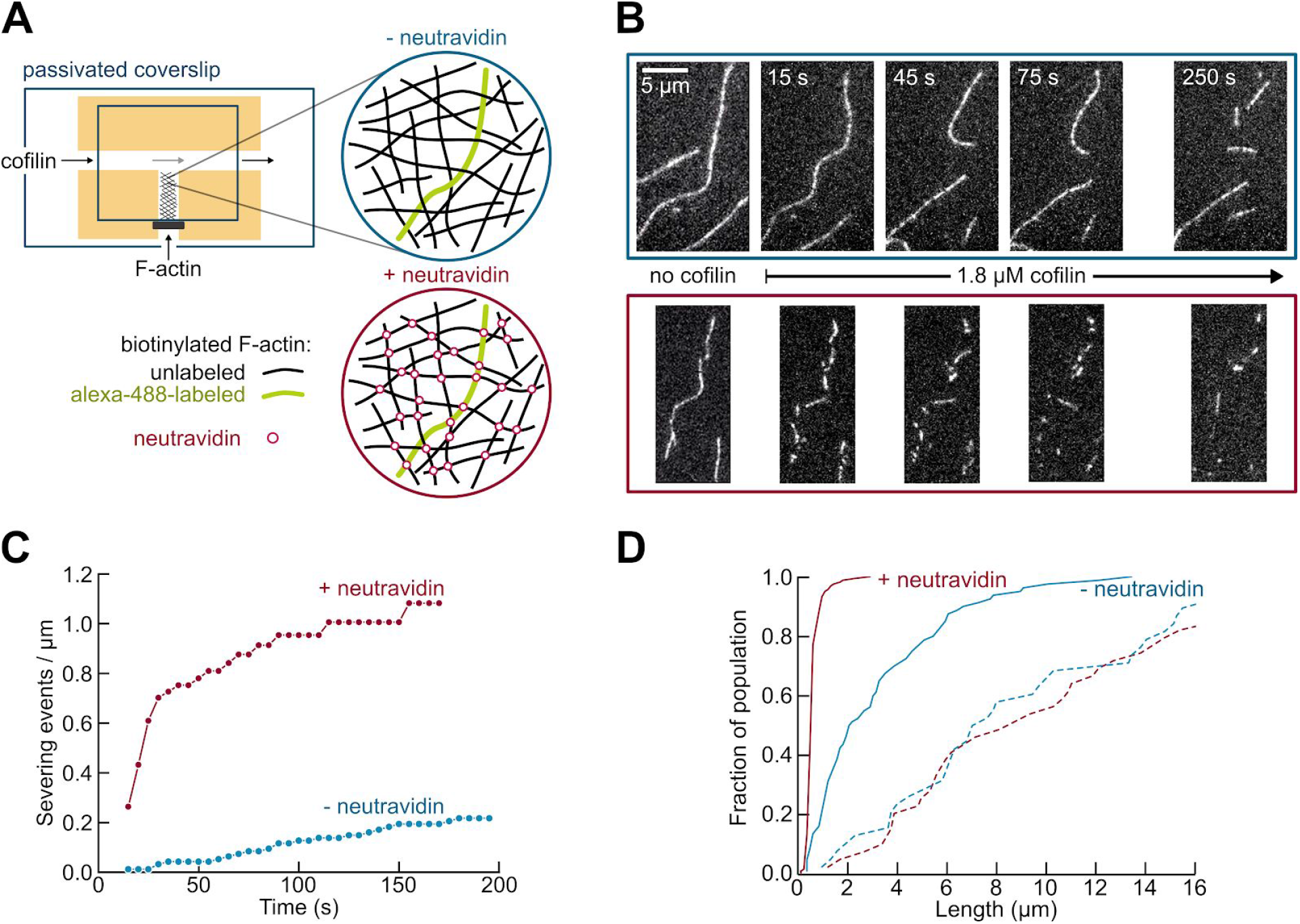
Enhanced severing of interconnected actin filament networks. (A) Experimental setup, using T-shaped chambers, seen from above. A solution of biotinylated F-actin, containing one fluorescently labeled filament for 20 unlabeled filaments, was first injected in the short channel. The filaments could either be left non-connected (top, blue) or be cross-linked by injecting neutravidin through the main channel (bottom, red). Cofilin was then injected in the main channel and F-actin severing was observed in a region near the channel junction. (B) Time-lapse of individual Alexa488-labeled filaments, within a meshwork of unlabeled filaments, prior to and after filling the main channel with 1.8 μM cofilin (at time t=0). Filaments are either non-connected (top, blue) or cross-linked by neutravidin (bottom, red). (C). Quantification of severing events, cumulated over time, for non-connected (blue, 10 filaments with a total initial length of 102 μm) and interconnected (red, 5 filaments with a total initial length of 60 μm) filaments, following the introduction of 1.8 μM cofilin in the main channel. (D) Cumulative length distributions (i.e., showing the fraction of a filament population having a length smaller than a given value) before exposure to cofilin (dashed lines) and 250 s after flowing 1.8 μM cofilin in the main channel (solid lines), for non-connected (blue, initial population of 38 filaments, final population of 80 observable filament fragments) and interconnected filaments (red, initial population of 39 filaments, final population of 162 observable filament fragments).

Methylcellulose was present in the buffer, in order to maintain the filaments close to the passivated surface at the bottom of the chamber, thus forming a dense, quasi bidimensional filament network. Different solutions could then be flown in the main channel of the chamber, and their components could diffuse into the chamber dead end, without mechanically perturbing the filament population. We first introduced either a neutravidin solution or buffer in the main channel, in order to either cross-link filaments or not. We then flew a solution of cofilin in the main channel, and observed its impact on the filaments. In each experiment, we monitored filaments in the same region of the chamber, 500 μm away from the channel junction.

Upon exposure to cofilin, the fates of the two filament populations were dramatically different, with the interconnected filaments experiencing far more severing (Fig. 5B-D). We have quantified the severing events in each population (Fig. 5C) and we can estimate that, shortly after flowing in cofilin, interconnected filaments severed more than 30-fold faster than non-connected filaments (with initial severing rates of approximately 0.001 and 0.035 event/μm/s for non-connected and interconnected filaments, respectively). On longer time scales, when the filaments were saturated by cofilin, many filaments of a few micrometers in length could be observed in the non-connected network, while only sub-micron fragments remained of the interconnected network (Fig. 5D). By creating new filament ends, severing also promotes the depolymerization of the filaments in the network: 250 s after flowing in cofilin, only 22% of the cross-linked F-actin remained visible (the rest being either fully depolymerized or in fragments too small to be detected) while 73% of the non-connected F-actin was still visible.

Compared to our single filament observations, severing appears to take place slower in our network experiment, possibly due to the diffusion and consumption of the finite cofilin pool in the closed, T-shaped microchamber, and to the presence of methylcellulose. Nonetheless, our experimental observations are in good quantitative agreement with the results of our simulations, which were based on our measured rates and our model (Fig. 4). From the observed filament density, and taking into account that there were 20 unlabeled filaments for every labeled filament, we could estimate that the cross-link density in our experiment was of the order of 1 μm^−1^. According to Fig. 4E we could thus expect that interconnected filaments would typically experience one severing event per micrometer, while most of the equivalent segments in non-connected filaments will saturate and not sever. This is indeed what we observed: after 200 s, interconnected filaments cumulated a bit more than one severing event per μm, while non-connected filaments were still several microns long on average (Fig. 5C-D).

### Implications for actin disassembly in cells

We show here that torsional stress and bending enhance filament severing by cofilin. These observations are consistent with early reports showing that, in the absence of cofilin, imposing a torque (31) or sharp bends (34) to actin filaments makes it easier to break them by applying tension. In our study, however, the torque is applied by cofilin itself as it binds to twist-constrained filaments, and the resulting torque is enough to dramatically increase the severing rate at the boundaries of cofilin domains. In cells, where filaments are typically interconnected and are not free to rotate, this mechanism is likely to play an important role. In particular, since its consequences are more drastic on densely connected filaments, it may modulate the disassembly of filament networks based on their crosslink density.

In cells, additional effects may come from the application of a torque to actin filaments by other factors. A recent theoretical study predicts that under-twisting an actin filament, beyond the maximum of approximately 5 rotations per micrometer that cofilin can induce on its own, would lead to an enhancement of cofilin dissociation, and that over-twisting would have a stronger effect (22). Torque may be applied to filaments as they are elongated by formins which are unable to rotate, and this formin-induced torque was recently reported to protect filaments from cofilin (21). These results suggest that formin-induced torque can reach higher values than the cofilin-induced torque we report here, and future studies will be needed to further explore the specificities of the different means to apply a torque to actin filaments.

The enhancement of severing by a cofilin-induced torque is a very general mechanism since all it requires is for filaments to be constrained in twist. This situation arises whenever filaments are anchored, or cross-linked, regardless of the molecular nature of the cross-links. Our results are sufficient to explain why severing by cofilin is enhanced when filaments are bundled by fascin (24). Consistently, when filaments are bundled by a crowding agent, without cross-links that would constrain their twist, no enhancement of severing is observed (35). Other factors, specific to different cross-linkers, may also modulate severing, in addition to the generation of a torque. For instance, severing may be further enhanced by cofilin discontinuities due to its competition with cross-linkers, or by local changes in stiffness due to the presence of cross-links (36). Bulky cross-linkers (37) or very tight filament packing may also alter cofilin’s access to the sides of the filaments.

The torque generated by the binding of cofilin onto twist-constrained filaments may also affect the binding of other regulatory proteins, such as tropomyosins (30, 38) or Aip1 (9, 17, 39), and thereby modulate the competition or the cooperative binding of these proteins. Cofilin-induced torque on interconnected filaments is thus likely to have consequences beyond the enhanced severing we report here, and may play an essential regulatory role in cells.

## Materials and methods

Detailed experimental procedures and data analysis used in this study are described in *SI Materials and Methods*.

### Proteins and buffer

Actin was purified from rabbit muscle and labeled on surface lysines with Alexa488-or Alexa568-succinimidyl ester. Recombinant human Cofilin-1 and ADF were expressed in E. coli and purified. All experiments were performed in F-buffer with 50 mM KCl (5 mM Tris-HCl pH 7.8, 50 mM KCl, 1 mM MgCl2, 0.2 mM EGTA, 0.2 mM ATP, 10 mM DTT and 1 mM DABCO).

### Microscopy experiments

Actin filaments were aged for at least 15 minutes after polymerizing in order to have fully ADP-actin, except for the experiments in Fig. 1. Microfluidics experiment were performed following the lines of our initial microfluidics experiments (40) where filaments were anchored by one end only to the coverslip surface, at the bottom of a flow chamber otherwise made of Poly Dimethyl Siloxane (PDMS).

### Data analysis

A Kaplan-Meier algorithm was applied to determine survival functions from the observation of individual events.

### Simulations

Numerical simulations followed a Gillespie algorithm, and programs were written in Python.

## ACKNOWLEDGEMENTS

We thank Martin Lenz for fruitful discussions, as well as Arnaud Echard, Nicolas Minc, Alexis Gautreau and Christophe Le Clainche for their critical reading of an early draft of this manuscript. We thank all members of the Romet/Jegou lab for their help, and especially Emiko Suzuki for initiating measurements on polarized fluorophore emission. We acknowledge funding from the Human Frontier Science Program (grant RGY0066 to G.R.-L.), the french ANR (grant Muscactin to G.R.-L.), the European Research Council (grant StG-679116 to A.J.) and Fondation ARC pour la Recherche sur le Cancer (postdoctoral fellowship to H.W.).

